# Tensions on the actin cytoskeleton and apical cell junctions in the *C. elegans* spermatheca are influenced by spermathecal anatomy, ovulation state and activation of myosin

**DOI:** 10.1101/2024.09.03.611016

**Authors:** Fereshteh Sadeghian, Noa W.F. Grooms, Samuel H. Chung, Erin J. Cram

## Abstract

Cells generate mechanical forces mainly through myosin motor activity on the actin cytoskeleton. In *C. elegans*, actomyosin stress fibers drive contractility of the smooth muscle-like cells of the spermatheca, a distensible, tube-shaped tissue in the hermaphrodite reproductive system and the site of oocyte fertilization. Stretching of the spermathecal cells by oocyte entry triggers activation of the small GTPase Rho. In this study, we asked how forces are distributed in vivo using the spermatheca, and explored how this tissue responds to alterations in myosin activity. Using laser ablation, we show that the basal actomyosin fibers are under tension in the occupied spermatheca. Reducing actomyosin contractility by depletion of the phospholipase C-ε/PLC-1 or non-muscle myosin II/NMY-1, leads to distended spermathecae occupied by one or more embryos, but does not alter tension on the basal actomyosin fibers. This suggests that much of the tension on the basal actin fibers in the occupied spermatheca is due to the presence of the embryo. However, activating myosin through depletion of the Rho GAP SPV-1 increases tension on the actomyosin fibers, consistent with earlier studies showing Rho drives spermathecal contractility. On the inner surface of the spermathecal tube, tension on the apical junctions is decreased by depletion of PLC-1 and NMY-1. Surprisingly, when basal contractility is increased through SPV-1 depletion, the tension on apical junctions also decreases, with the most significant effect on the junctions aligned in perpendicular to the axis of the spermatheca. This suggests tension on the outer basal surface may compress the apical side, and suggests the three-dimensional shape of the spermatheca plays a role in force distribution and contractility during ovulation.

## Introduction

Mechanical forces regulate essential cellular processes such as adhesion, signaling, proliferation, and differentiation. Throughout morphogenesis, mechanical forces alter cell shapes (1). Cells generate mechanical forces predominantly through the action of the motor protein myosin on the actin cytoskeleton (1,2). The actin cytoskeleton is a dynamic network formed by the polymerization of globular actin monomers (G-actin) into helical filaments (F-actin). Actin networks are crucial for various cellular processes such as cell migration, tissue morphogenesis, wound healing, and the adaptation of cells and tissues to physical stress (3). Stress fibers are contractile actomyosin bundles, commonly found in cells experiencing physical stress (3). Cells use actomyosin stress fibers to exert traction forces against focal adhesions, integrin-based adhesive structures that link the cytoskeleton to the extracellular matrix (ECM) (4).

To study coordinated actomyosin contractility in an intact context, we use the *Caenorhabditis elegans* reproductive system as a model. The hermaphrodite gonad consists of two symmetrical gonad arms containing germ cells and oocytes surrounded by sheath cells, two spermathecae, and a common uterus (Figure 1A) (5). The spermatheca, which stores sperm and serves as the site of fertilization in the hermaphrodite, is a contractile tube comprising 24 smooth muscle-like cells. The spermathecal-uterine (sp-ut) valve is a contractile sphincter that connects the spermatheca to the uterus. The apical surface of the spermatheca faces the oocyte. Upon the entry of the oocyte (Figure 1B), immediate fertilization occurs, initiating the formation of the eggshell. Subsequently, the spermatheca undergoes coordinated contractions to propel the embryo through the sp-ut valve and into the uterus (6).

**Figure 1:**
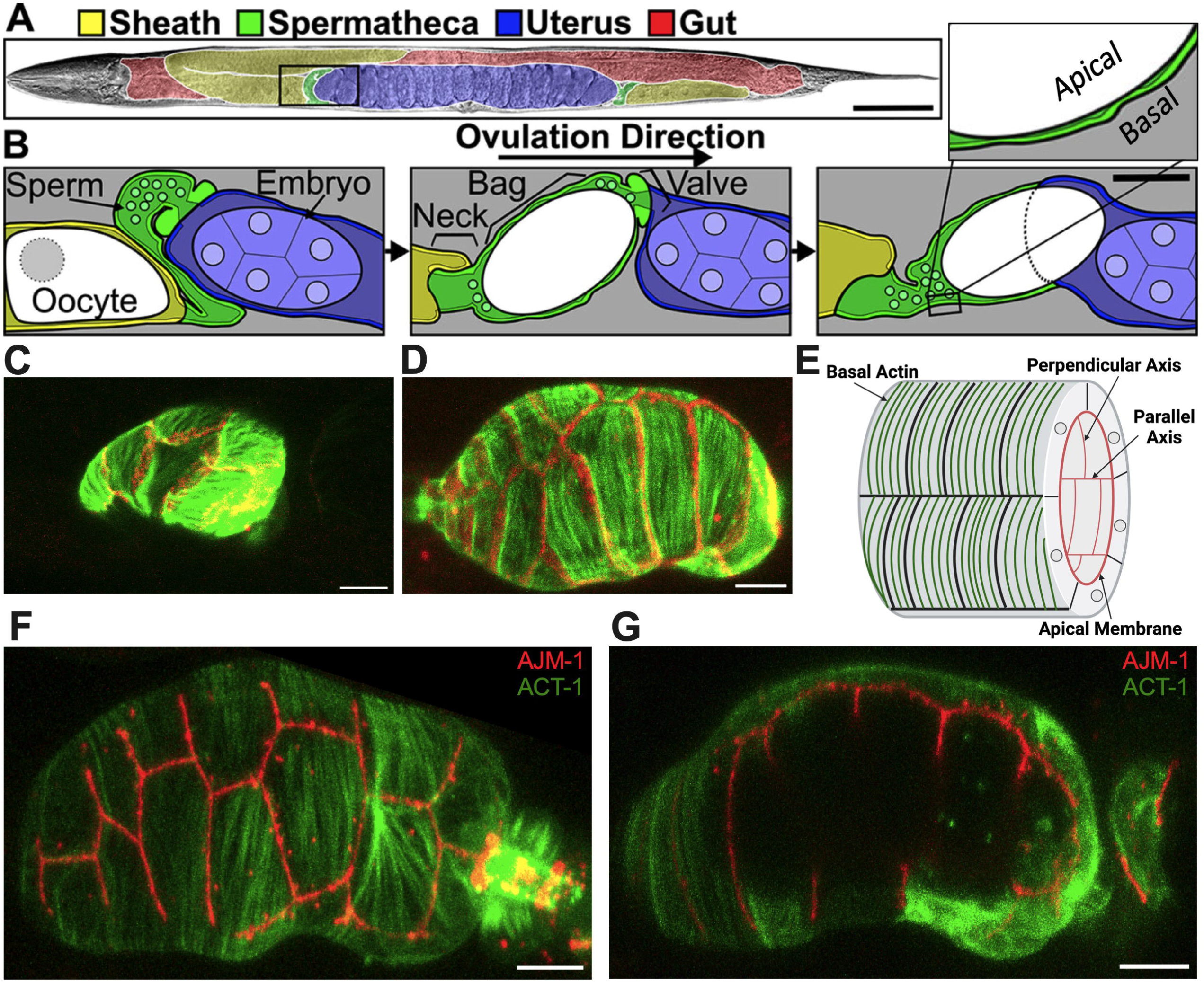
Structure of the spermatheca. (A) Brightfield image of an adult hermaphrodite false colored to indicate the sheath cells (yellow), spermathecae (green), uterus (blue), and gut (red). (B) Diagram of the area indicated by a black box in A during an ovulation. First panel: sheath cell contractions begin to push the proximal oocyte (white). Second panel: sheath contractions force the oocyte into the spermatheca, where it is fertilized. Used by permission (7). Third panel: the spermathecal-uterine valve opens as the spermathecal bag contracts to expel the fertilized embryo into the uterus. Insert in B shows a magnified cross-section of the spermatheca indicating that the apical surface faces the lumen. Scale bars, 50 μm in A and 20 μm in B. (C) Unoccupied spermatheca showing green actin ACT-1::GFP and red gap junction INX-12::mApple expression before oocyte entry. (D) Occupied spermatheca showing aligned green actin and red gap junctions during ovulation. (E) Diagram of the spermatheca indicating basal actin in green and apical membrane in red, indicating the membranes parallel and perpendicular to the axis of spermatheca. Created with Biorender.com. (F) Basal Actin/ACT-1::GFP and Apical junction/AJM-1::RFP expression in spermatheca. (G) A cross section of the same spermatheca in (F) showing of green basal actin on the exterior surface and red apical membrane on the interior surface. Scale bar, 20 μm.

Stress fiber-like bundles of actin and myosin drive spermathecal contractility (3). These bundles are on the outer, basal surface of the spermathecal cells and in most spermathecal cells are aligned perpendicular to the axis of the spermatheca (3). The inner, luminal sides of the cells are connected by apical cell-cell junctions (7). Prior to oocyte entry, the spermatheca is relaxed and compact (Figure 1C); during ovulation cells are stretched by the presence of oocyte (Figure 1D). The diagram in Figure 1E depicts the spermathecal anatomy with basal actin fibers and the interior apical junctions. Figure 1F gives the exterior view, and Figure 1G gives the interior view of the occupied spermatheca, with red apical junctions and green basal actin.

The contraction of actomyosin fibers in the spermatheca is regulated by highly conserved pathways similar to non-muscle and smooth muscle cell cascades (5). Briefly, activation of the phospholipase PLC,/PLC-1 leads to cleavage of the membrane lipid PIP_2_, resulting in the production of IP_3_ (5). Binding to the IP_3_ receptor triggers the release of Ca^2+^ from the endoplasmic reticulum and subsequent activation of the myosin light chain kinase/MLCK-1 (8). In a parallel pathway, oocyte entry causes displacement of the RhoGAP SPV-1, activating RhoA/RHO-1 and ROCK/LET-502 (9). This activation leads to the phosphorylation and activation of non-muscle myosin II/NMY-1, resulting in the contraction of actomyosin fibers (5). Activation and coordination of both pathways is necessary for embryos to successfully pass through the spermatheca. SPV-1, a BAR-domain-containing Rho GAP, functions as a tension sensor in the spermatheca, localizing to low-tension membranes where it maintains Rho/RHO-1 in an inactive state (10). Prior to the first ovulation, the spermathecal actin forms a diffuse, weblike-network (3,8). Activation of myosin pulls the actomyosin fibers into aligned, parallel arrays (8). For productive contractility, the fibers must be anchored. Filamins form a resilient three-dimensional network of the actin cytoskeleton and establish links between the cytoskeleton and transmembrane proteins, including integrins (11). Filamin/FLN-1 is essential for anchoring actin bundles at the basal surface of the spermatheca, and preventing the aggregation of actin and myosin near the nucleus (6).

Observation of the actin cytoskeleton during ovulation led us to the hypothesis that the parallel arrays of actomyosin fibers are under tension and that activating myosin would lead to an increase in tension, productive contraction of the spermatheca, and expulsion of the embryo into the uterus. Exit of the embryo from the spermatheca requires coordinated contraction of the distal spermathecal cells (near the entry neck) and dilation of the sp-ut valve (9). We wondered whether mechanical tension, or increased tension, on the fibers or the cell-cell junctions might provide temporal or positional information to regulate spermathecal contractility or embryo exit. To explore these ideas, we needed to first determine whether the fibers and/or cell junctions are under tension and, if so, whether the perpendicular or parallel junctions might be under higher load.

The mechanical characteristics of actomyosin stress fibers, epithelial cell–cell junctions, and other cellular structures can be explored using laser ablation (12). Femtosecond laser ablation enables minimally invasive disruption of cellular structures with sub-micrometer resolution (13,14). Laser ablation has been used for studying the viscoelastic response of myofibrils of iPSC-derived cardiomyocyte (12), force transmission between apical and basal nucleus by actomyosin dynamics in *Drosophila* (15), and tension in cell-cell boundaries in *Drosophila* follicular epithelium (16) and the amnioserosa (17).

In this study, we employed imaging, laser ablation, and quantitative image analysis to characterize the tensions in the spermatheca. We demonstrate that decreasing actomyosin contractility through depletion of phospholipase C-ε/PLC-1, or non-muscle myosin II/NMY-1, did not alter tension on the basal actomyosin fibers, suggesting much of the tension is coming from the presence of the embryo, at least in the central spermathecal cells and at the timepoint selected. Activating myosin through depletion of the Rho GAP SPV-1 does increase tension on the basal actomyosin fibers. Conversely, increasing contractility through SPV-1 depletion decreased tension on the apical junctions, suggesting that tension on the outer, basal surface of this tubular structure might compress the apical side.

## Materials and methods

### Strains and culture

Nematodes were grown on nematode growth media (NGM) (0.107 M NaCl, 0.25% wt/vol Peptone (Fischer Science Education), 1.7% wt/vol BD Bacto-Agar (FisherScientific), 0.5% Nystatin (Sigma), 0.1 mM CaCl_2_, 0.1 mM MgSO_4_, 0.5% wt/vol cholesterol, 2.5 mM KPO_4_) and seeded with *E. coli* OP50 using standard *C. elegans* techniques. Nematodes were cultured at 23°C for 52–72 hr (5). Lines expressing ACT-1::GFP (UN1502) and AJM-1::GFP (UN2140) were previously generated by injecting plasmids and following integration (6,18). Strains used to show the basal and apical side in Figure 1 were generated by crossing ACT-1::GFP and INX-12::mApple (UN1617), ACT-1::GFP and AJM-1::RFP (UN1936).

### RNA interference

The RNAi protocol was performed essentially as described in Timmons et al (1998). HT115(DE3) bacteria (RNAi bacteria) transformed with a dsRNA expression construct of interest was grown overnight in Luria Broth (LB) supplemented with 40 μg/ml ampicillin and seeded (150 μl) on NGM plates supplemented with 25 μg/ml carbenicillin and 1 mM disopropylthio-β-galactoside (IPTG). Seeded plates were left for 24–72 hours at room temperature (RT) to induce dsRNA expression. Empty pPD129.36 vector (“Control RNAi”) was used as a negative control in all RNAi experiments (5). Embryos from gravid adults were collected using an alkaline hypochlorite solution as described by Hope (1999) and washed three times in M9 buffer (22 mM KH_2_PO_4_, 42 mM NaHPO_4_, 86 mM NaCl, and 1 mM MgSO_4_) (‘egg prep’). Clean embryos were transferred to supplemented IPTG plates seeded with HT115(DE3) bacteria expressing dsRNA of interest and left to incubate at 23°C for 50–56 hours depending on the experiment (5).

### Population assay

Embryos collected via an ‘egg prep’ as previously described were plated on supplemented NGM-IPTG plates seeded with RNAi bacterial clones of interest. Plates were incubated at 23°C for 54– 56 hours or until animals reached adulthood. Upon adulthood nematodes were killed in a drop of 0.08 M sodium azide (NaAz) and mounted on 2% agarose pads to be visualized using a 60x oil-immersion objective with a Nikon Eclipse 80i epifluorescence microscope equipped with a Spot RT3 CCD camera (Diagnostic instruments; Sterling Heights, MI, USA). Animals were scored for the presence or absence of an embryo in the spermatheca.

### Fluorescence microscopy image acquisition and processing

For time-lapse imaging of ovulation and observations of live or fixed animals, partially synchronized populations were obtained by egg prep and animals were grown at 23°C for ∼54 h, around the time of the first ovulation. Live animals were immobilized with 1.85% formaldehyde in PBS for 25 min at room temperature, washed twice in PBS and mounted on a 3% agarose pad. Confocal microscopy was performed on an LSM 710 confocal microscope (Zeiss) equipped with Zen software (Zeiss) using a Plan-Apochromat 63×/1.40 oil DIC M27 objective. A 488-nm laser was used for GFP and DyLight 488, and a 561 nm laser was used for RFP and TexasRed. (18).

### Laser Severing

A Yb-doped diode-pumped solid-state laser (Spectra-Physics Spirit-1040-4W) outputs 1040-nm 400-fs pulses with a variable repetition rate, set to 1 kHz. We focused laser pulses with a Nikon CFI Plan Apo Lambda 1.4 NA 60x oil immersion objective. We set the laser power to 150 µW (14).

### Statistics

Either a Fishers exact t-test (two dimensional x2 analysis) or a one-way ANOVA with a multiple comparison Tukey’s test were conducted using GraphPad Prism on total. For population assays, statistics were performed using the total number of unoccupied spermathecae compared with the sum all other phenotypes. N is the total number of spermathecae counted. Fisher’s exact t-test were used for population assays. Stars designate statistical significance (**** p<0.0001, *** p<0.005, ** p<0.01, * p<0.05). (5). Non-linear regression analysis was used to show correlation between the length of retraction and the length or width of the spermatheca.

## Results

### Basal actomyosin fibers in the spermatheca are under tension

The spermatheca expands upon oocyte entry and contracts when the embryo is expelled into the uterus (Figure 1). We have shown that oocyte entry stimulates the rearrangement of the cytoskeleton into parallel, aligned actomyosin bundles on the basal surface of the spermathecal (6) (Figure 1). As described above, spermathecal contractility is regulated by two parallel pathways, one which requires the phospholipase PLC-1/PLC, and the other that requires the small GTPase RHO-1/Rho (5). We have shown previously that *plc-1* and *nmy-1* are required for successful transit of the embryo through the spermatheca (5,6). Failed transits lead to an increase in the percentage of spermathecae occupied by an embryo. To establish that the conditions to be used in our laser severing experiments do disrupt spermathecal contractility, and to confirm these prior results, we determined the spermathecal occupancy in each RNAi condition and found that, as expected, depletion of *plc-1* and *nmy-1* lead to 100% and 90% occupancy compared to control (empty vector), respectively (Figure 2A).

**Figure 2:**
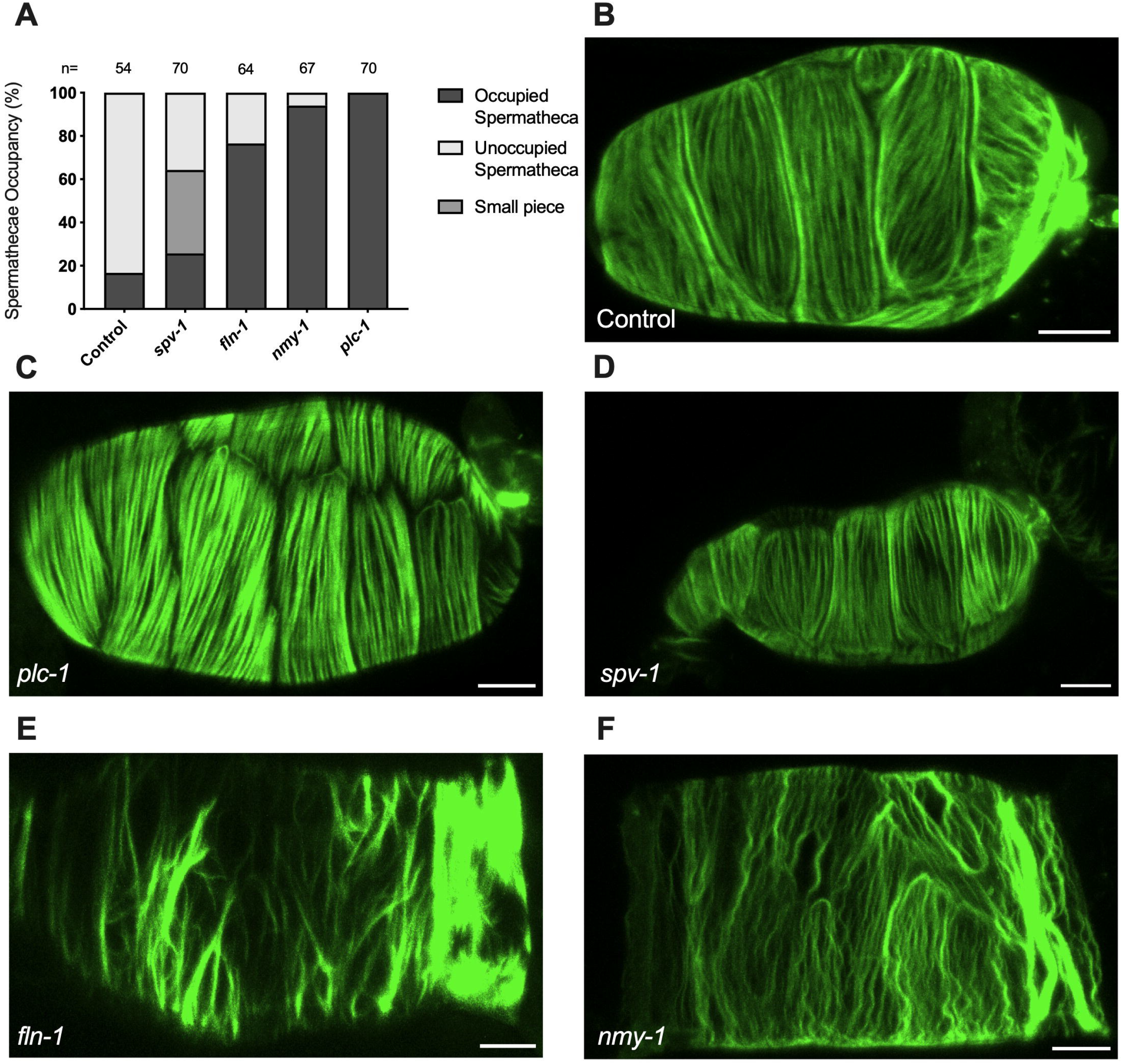
Phospholipase and Rho signaling regulate contractility in spermatheca. (A) Spermathecal occupancy in wild type ACT-1::GFP animals cultivated on control, *plc-1* RNAi, *spv-1* RNAi, *fln-1* RNAi, and *nmy-1* RNAi. N represents the total number of spermathecae counted. Actin phenotype of wild type animals (ACT-1::GFP) grown on control (B), *plc-1* RNAi (C), *spv-1* RNAi (D), *fln-1* RNAi (E), and *nmy-1* RNAi (F). Scale bar, 20 μm.

We also determined whether the depletion of *plc-1* and *nmy-1* led to the expected effects on the actin cytoskeleton (Figure 2). As expected, PLC-1 plays a minor role in the formation of the parallel fiber bundles (Figure 2C) (18) whereas *nmy-1*/myosin depletion results in the failure to form parallel, aligned fiber bundles (Figure 2F). FLN-1/filamin is essential for the development and stabilization of a regular arrangement of parallel, contractile actomyosin fibers in the spermatheca (6). As expected, depletion of FLN-1 disrupts anchoring of fibers in the actin network, leading to an inability to contract (Figure 2E). This defect also results in the trapping of oocytes in the spermatheca (6).

We reasoned that the presence of the embryo exerts tension on the fibers, and that activation of myosin would further increase the tension on the fibers, eventually leading to expulsion of the embryo from the spermatheca. To explore these ideas, we used laser severing to cut basal actin fibers in animals expressing ACT-1::GFP in the spermatheca. The movement of GFP-labeled actin fibers was tracked for up to 30 seconds after ablation from the point of surgery (Supplemental Movies S1 and 2). The first 10 seconds of each retraction is shown in Figure 3D-G, during which time the retraction plateaued in all samples.

**Figure 3:**
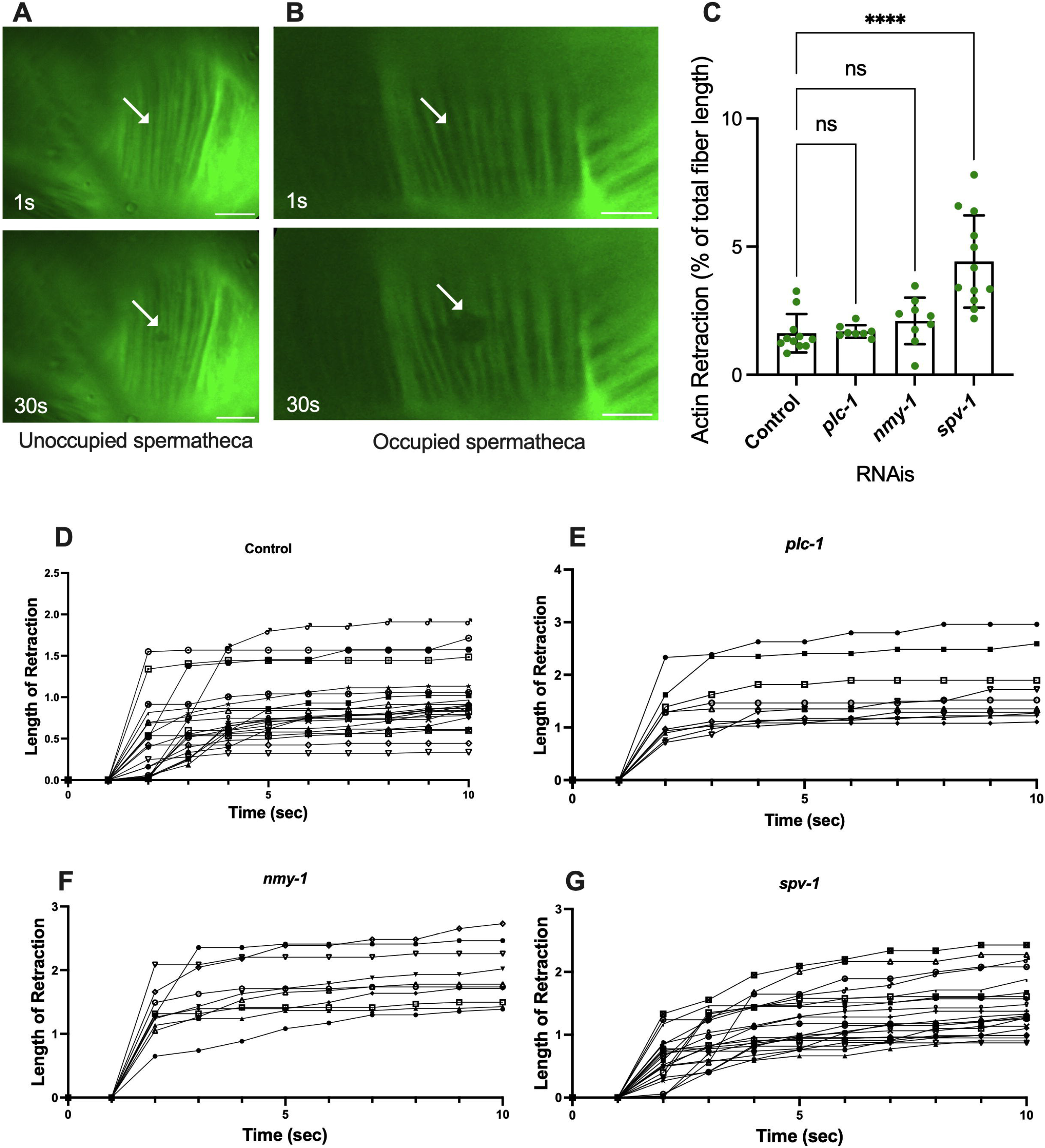
Basal fibers are under tension. Laser ablation of ACT-1::GFP expressing animals treated with control RNAi, in the unoccupied spermatheca (A), and occupied spermatheca (B) before and 30s after surgery, (see *plc-1, nmy-1*, and *spv-1* RNAis in Figure S1). Actin fibers severed using a femtosecond laser. (C) Total retraction of actin fibers in control (empty vector), *plc-1, nmy-1*, and *spv-1* RNAi. Data indicates higher tension in *spv-1* RNAi treated animals with no significant differences between empty, *plc-1*, and *nmy-1* RNAi treatments. Retraction of control compared to other conditions was conducted using Ordinary one-way ANOVA, *\*\*\*\*p < .0001*. (D-G) Analysis of length of retraction to the length of the fiber shows tension release of actin fibers after laser ablation in ACT-1::GFP expressing animals treated with control (D), *plc-1* RNAi (E), *nmy-1* RNAi (F), and *spv-1* RNAi (G). Scale bar, 10 μm.

The actin fibers in the central cells of the spermatheca are aligned circumferentially to the axis of the spermatheca (as shown in Figure 1). All incisions were executed perpendicular to the axis of the spermatheca. Fiber retraction was normalized to the length of the fiber in all conditions (Figure 3C and Figure 5C-E). The length of retraction after ablation was measured (Supplemental Figure S2A (actin) and Supplemental Figure S2B-D (membrane)). Then, because the size of spermatheca was different for each condition, the length of the retraction was divided by the length of the corresponding fiber before incision. In some laser ablations, more than one actin fiber was severed in an incision, in this case, the center expansion of the lesion was measured.

To confirm that the fibers were severed, rather than photobleached, we tracked fluorescence recovery after laser ablation, and verified no recovery of the GFP signal in the 30 sec following the incision in analysis. To determine whether the fibers depolymerized, rather than retracting, we investigated the length of initial retraction and the following retraction of the severed fibers throughout the time (Figure 3). In four animals, the ablated actin fibers continued to shorten at a slower rate after the initial retraction. These were excluded from further analysis. All retraction data is shown in supplementary Figure S2.

To determine whether fiber retraction is influenced by the presence of an oocyte in the spermatheca, we severed fibers in occupied and unoccupied spermathecae. This was challenging in the unoccupied spermatheca, given the folded morphology of the unoccupied tissue (Figure 3A). However, no actin fiber retraction was observed in the empty spermathecae (Figure 3A). This is in contrast to the significant retraction observed in occupied spermatheca under the same severing conditions (Figure 3B). This suggests that retraction depends on the presence of the oocyte in the spermatheca, and that the basal fibers are under tension in the occupied spermatheca.

### SPV-1 depletion increases tension on basal actomyosin fibers in spermatheca

We hypothesized that decreasing contractility would decrease the tension on the actin fibers, and increasing contractility would increase tension. To test this, we treated the ACT-1::GFP animals with control RNAi, *plc-1* RNAi, *nmy-1* RNAi to decrease contractility, and severed actin fibers in occupied spermathecae. To our surprise, inhibiting actomyosin contractility did not affect the tension on the fibers. Following the incision, the actin retracted to the same extent as controls in *plc-1* and *nmy-1* depleted cells (Figure 3C). Because reducing actomyosin contractility does not affect the measured tensions, this suggests most of the tension may come from the presence of the embryo at least in the central cells of the spermatheca and at the timepoint chosen.

We next asked if increasing contractility would increase the tension on the actin fibers. The RhoGAP SPV-1 inactivates RHO-1 leading to decreased contractility. When *spv-1* is depleted by RNAi, the spermatheca is hypercontractile, often severing the oocytes upon entry or exit (9). As expected, actin fiber ablation in *spv-1* depleted animals shows an increase in fiber retraction, compared to controls (Figure 3C). We attempted to additionally increase tension in the spermatheca through depletion of the myosin phosphatase MEL-11. However, *mel-11* depletion resulted in spermathecal explosion due to excessive tension on fibers (8). The *spv-1* depletion result suggests activating myosin does increase tension on the basal actomyosin fibers.

### Distention of the spermatheca does not significantly affect fiber tension

We observed that depletion of PLC-1 and NMY-1 leads to the entrapment of more than one embryo, distending the spermatheca. Conversely, spermathecae treated with *spv-1* RNAi are smaller than wildtype and often contain small pieces of oocytes due to increased contractility and severing of oocytes (Figure 2D). To quantify these effects, we measured the length and width of ACT-1::GFP expressing spermathecae in control and RNAi depleted animals, and saw significant increases in both length and width in the *plc-1* and *nmy-1* RNAi and significant decreases in length and width in *spv-1* RNAi treated animals compared to controls (Figure 4A and B). We next asked if there was a relationship between degree of distention of the spermatheca and tension on the actin fibers. Due to the imaging setup, it was not possible to measure the length or width of the spermatheca in the same animals in which fibers were severed. Therefore, we averaged the length and width of spermatheca from the data shown in Figure 4 (A and B), and plotted those averages against the average retraction of the actin fibers seen in the laser ablation experiments.

**Figure 4:**
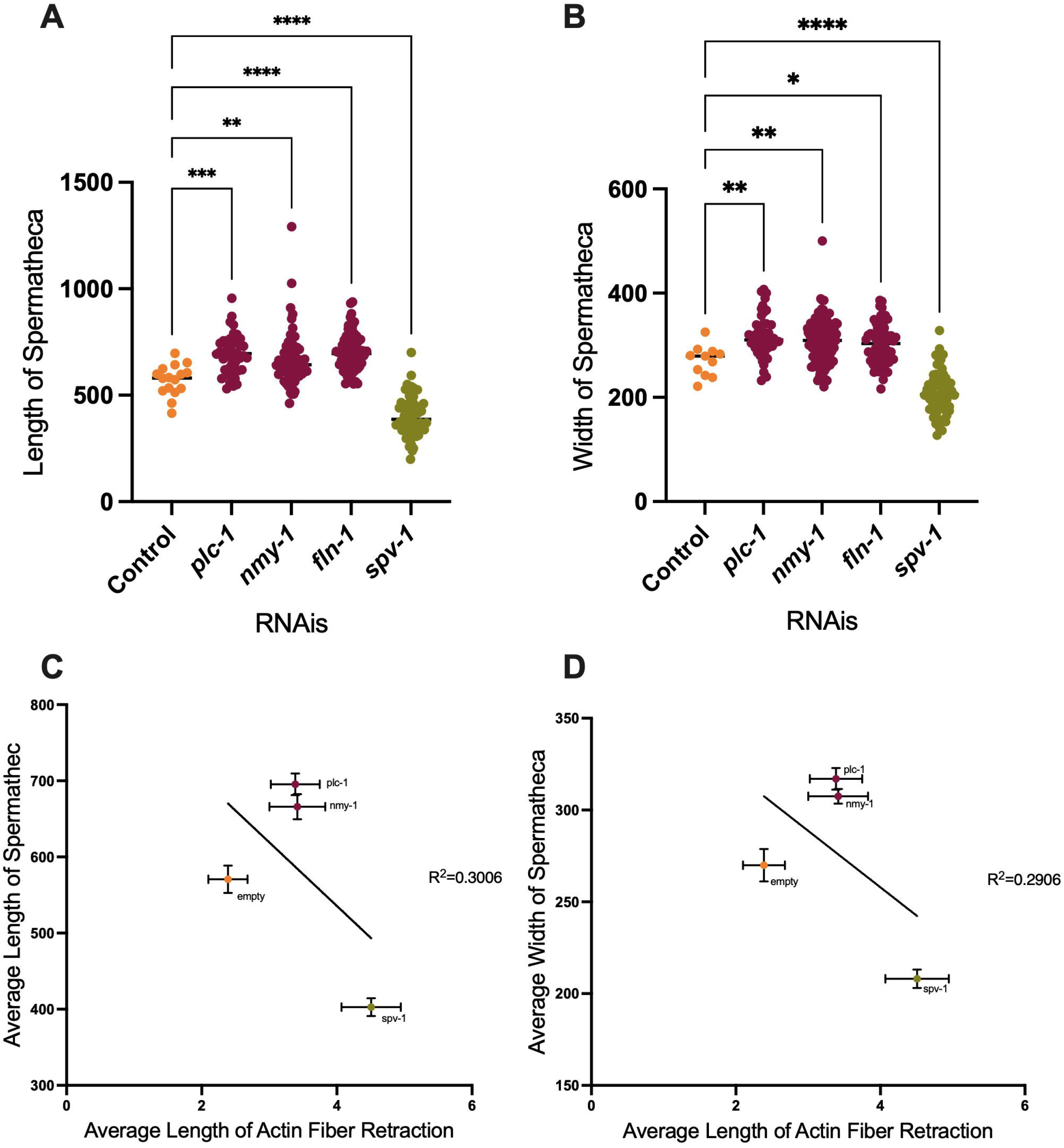
Depletion of contractility regulators affects spermathecal dimensions. Length and width of spermatheca of ACT-1::GFP (A and B). (C-D) Correlation between length (C) and width (D) of spermatheca to the length of retraction. Average of actin retraction to the average of length and width of spermatheca is shown for control, *plc-1* RNAi, *nmy-1* RNAi, and *spv-1* RNAi conditions. Non-linear regression analysis was used to assess correlation. Length of control (empty vector) compared to other conditions analyzed by using Ordinary one-way ANOVA, *\*\*\*\*p < .0001*. Scale bar 20 μm.

**Figure 5:**
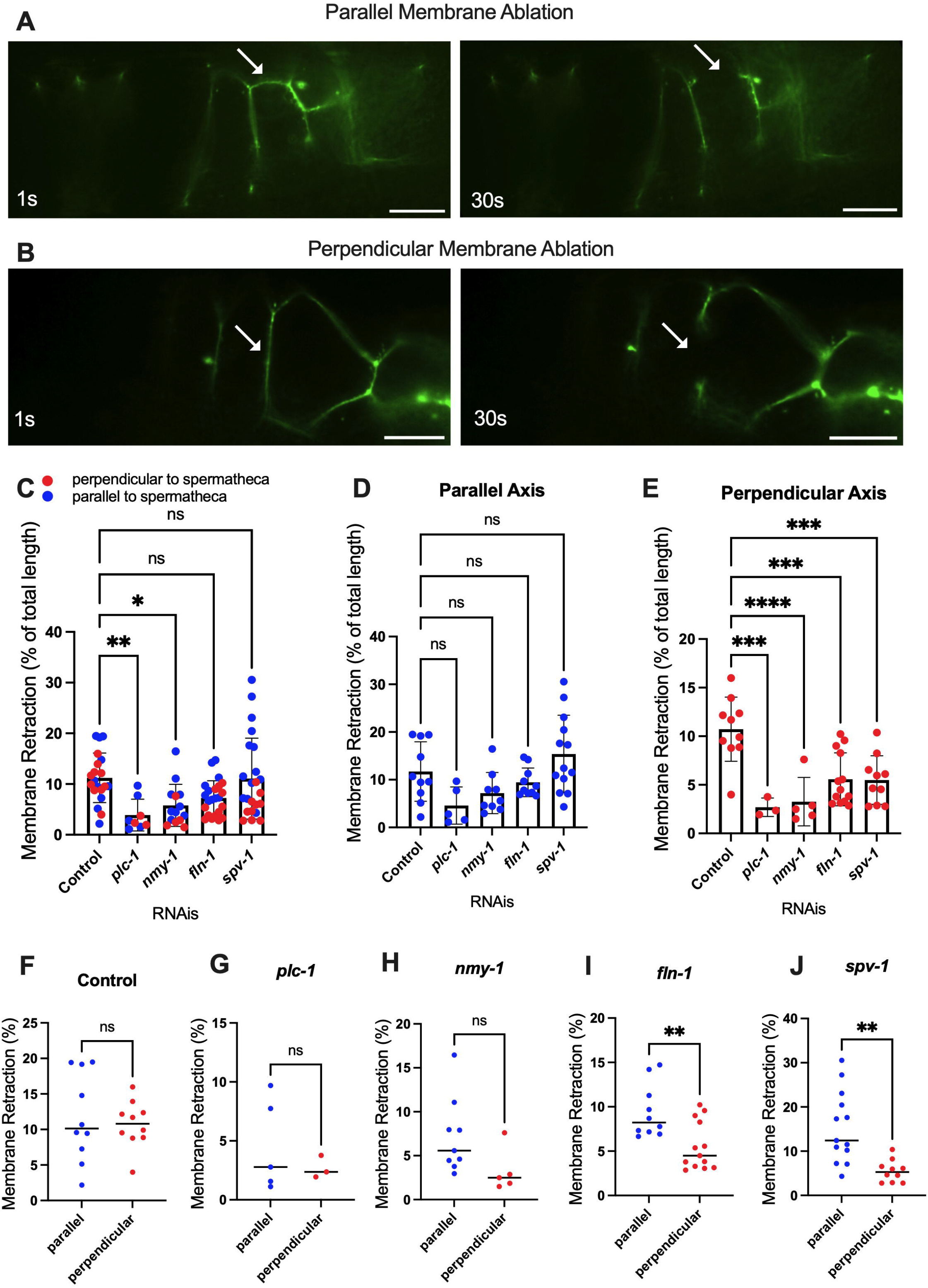
Apical junctions are under tension. Laser ablation of AJM-1::GFP expressing animals treated with control (empty vector) RNAi (A,B) before and 30 sec after surgery (see *plc-1* RNAi, *nmy-1* RNAi, *fln-1* RNAi, and *spv-1* RNAi in Figure S3 and S4). Parallel (A) and perpendicular (B) membranes were severed using a femtosecond laser. (C-E) Percentage of the total retraction normalized to the length of the membrane indicates changes in tension on parallel (D) and perpendicular (E) membranes with *plc-1* RNAi, *nmy-1* RNAi, *fln-1* RNAi, and *spv-1* RNAi treatments compared to control (F-J). Percentage of the total retraction normalized to the length of the membrane for each condition individually, comparing parallel and perpendicular membranes in control (F), *plc-1* RNAi (G), *nmy-1* RNAi (H), *fln-1* RNAi (I), and *spv-1* RNAi (J). Retraction of control (empty vector) compared to other conditions using Ordinary one-way ANOVA, *\*\*\*\*p < .0001*. Scale bar, 10 μm.

When PLC-1 and NMY-1 are depleted, the spermatheca are distended, and on average, there is slightly more tension on actin fibers, compared to the control condition. However, the fiber tensions in these RNAis are not significantly different from controls. In contrast, in *spv-1* RNAi treated animals with smaller spermatheca, the tension on actin is significantly increased (Figure 4C and D). These results suggest that the tension on the fibers does not scale with degree of distension of the tissue.

### Apical cell junctions/membranes are under tension

Entry of the oocyte triggers actin rearrangement and actomyosin contractility (9). We speculated that tension on the cells of the spermatheca might also convey temporal or spatial information to inform the directional contraction that expels the fertilized embryo from the spermatheca. The circumferential actin fibers in the central spermathecal cells are not suitable for conveying lateral information. To address the question of whether mechanical information might be transmitted from cell to cell in the spermatheca, we first asked if the apical membranes and apical junction complexes might be under tension, and whether the tension differed in the parallel or perpendicular cell membranes.

To investigate tension along both axes of the cell membrane, we performed apical junction ablation parallel and perpendicular to the axis of the spermatheca in separate cells (Figure 5A and B), (Supplemental Movies S3 and 4). We used laser severing to target the apical plasma membrane labeled with the apical junction marker AJM-1::GFP. In control conditions, the length of membrane retraction was similar in fibers parallel and perpendicular to the spermatheca (Figure 5C-F). However, when myosin contractility was reduced using *plc-1, nmy-1* and *fln-1* RNAi, we observed decreased membrane retraction in both parallel and perpendicular conditions compared to control, suggesting a decrease in membrane tension (Figure 5C). Most of this effect is the result of decreased membrane tension on fibers perpendicular to the axis of the spermatheca (compare Figure 5D and 5E). When we increased tension by depleting *spv-1*, we saw no overall change in the tension on apical membranes compared to control (Figure 5C), suggesting that most of the effect of SPV-1 is on the basal actomyosin fibers. However, we did observe a significant decrease in apical junction tension on perpendicular, but not parallel, apical membranes (Figure 5D and 5E). We also compared tensions on perpendicular versus parallel membranes for each gene depletion. No difference in tension was observed in control or the *plc-1* and *nmy-1* RNAi conditions (Figure 5F-H). Depletion of *fln-1*, which disrupts both the actin cytoskeleton and actomyosin contractility (6) resulted in a significant decrease in tension on perpendicular membranes (Figure 5I). Depletion of *spv-1* also resulted in a significant decrease in tension on perpendicular membranes (Figure 5J). The *spv-1* results suggest increased contraction on the basal side of the spermatheca may reduce apical tension by compressing the tissue.

## Discussion

In the spermatheca, actin and myosin are bundled into basal stress fiber-like actomyosin bundles. These bundles form during the initial ovulation and drive tissue contractility. We found that these fibers are under significant tension, influenced by both the presence of the oocyte and the activity of contractile proteins like SPV-1. Using laser ablation and RNA interference (RNAi), we demonstrate that decreasing contractility via depletion of *nmy-1*/myosin and *plc-1*/phospholipase C did not significantly affect fiber retraction, which suggests most of the tension on the fibers comes from the presence of the oocyte. However, increasing contractility by depleting SPV-1, which inactivates RHO-1, led to increased fiber retraction. This suggests that although the presence of the embryo strains the actomyosin fibers, activation of the Rho pathway can lead to further increases in contractility.

The apical, lumen-facing surface of the spermathecal cells are connected with apical junctions. Apical junctions are also under tension in the occupied spermatheca. Reducing contractility, particularly through loss of FLN-1/filamin, decreases this tension. Loss of FLN-1 may have a stronger effect than loss of PLC-1 or NMY-1 due to its role in maintaining the overall structure of the cell and the basal and apical actin cytoskeletons (6) in addition to regulating contractility per se (11). Perpendicular membranes are most strongly affected. This suggests that presence of the oocyte strains the width, or girth, of the spermatheca more than the length. This makes sense given the geometry of the spermatheca, which is a fairly narrow tube. We hypothesized that increasing contractility would increase tension on apical junctions as the tissue was pulled tightly against the egg. However, we found that when actomyosin contractility is increased through the depletion of SPV-1, apical tension is reduced. This suggests that increasing tension on the basal (outer) side of this biological tube compresses the inner the apical side.

Laser ablation studies in other systems show actomyosin network and dynamics play a key role in the mechanical properties and morphologies of living cells (1) (19) (20). Our work suggests the correct amount of tension is needed for tissue function. Similarly, in *C. elegans*, laser severing has been used to show CAP-1/actin capping protein limits myosin activity and cortical tension in the rachis of the hermaphrodite gonad (21). In *Drosophila* embryos, manipulation of myosin phosphorylation and actin polymerization give rise to changes in stiffness and contractility throughout the tissue, highlighting the importance of the cytoskeletal dynamics and transmitting contractile forces in developing tissues (1). A laser severing study of wound healing in *Drosophila* embryos shows the necessity of myosin dynamics for stability and wound repair (22). Cell and tissue shape have a significant influence on tensions. Similar to our observations, a study of tension and viscoelasticity of the sarcomere in single cells and in monolayers shows that the length and orientation of the stress fibers affect the retraction of the fibers (4).

This study sheds light on the temporal dynamics of contractility and tension, the role of various cytoskeletal components, and how mechanical signals are integrated across the tissue to coordinate the complex process of oocyte transit. These findings enhance our understanding of the mechanical properties of reproductive tissues and the interplay between cellular structures and contractile proteins in regulating these properties.

## Acknowledgements

We thank members of the Cram and Apfeld labs for helpful discussions. Some strains were provided by the CGC, which is funded by NIH Office of Research Infrastructure Programs (P40 OD010440). This work was supported by a grant from the National Institutes of Health National Institute of General Medical Sciences (GM110268) to E.J.C.

