## Supplementary figures and images for "Tensions on the actin cytoskeleton and apical cell junctions in the *C. elegans* spermatheca are influenced by spermathecal anatomy, ovulation state and activation of myosin"

### Supplemental Figure 1

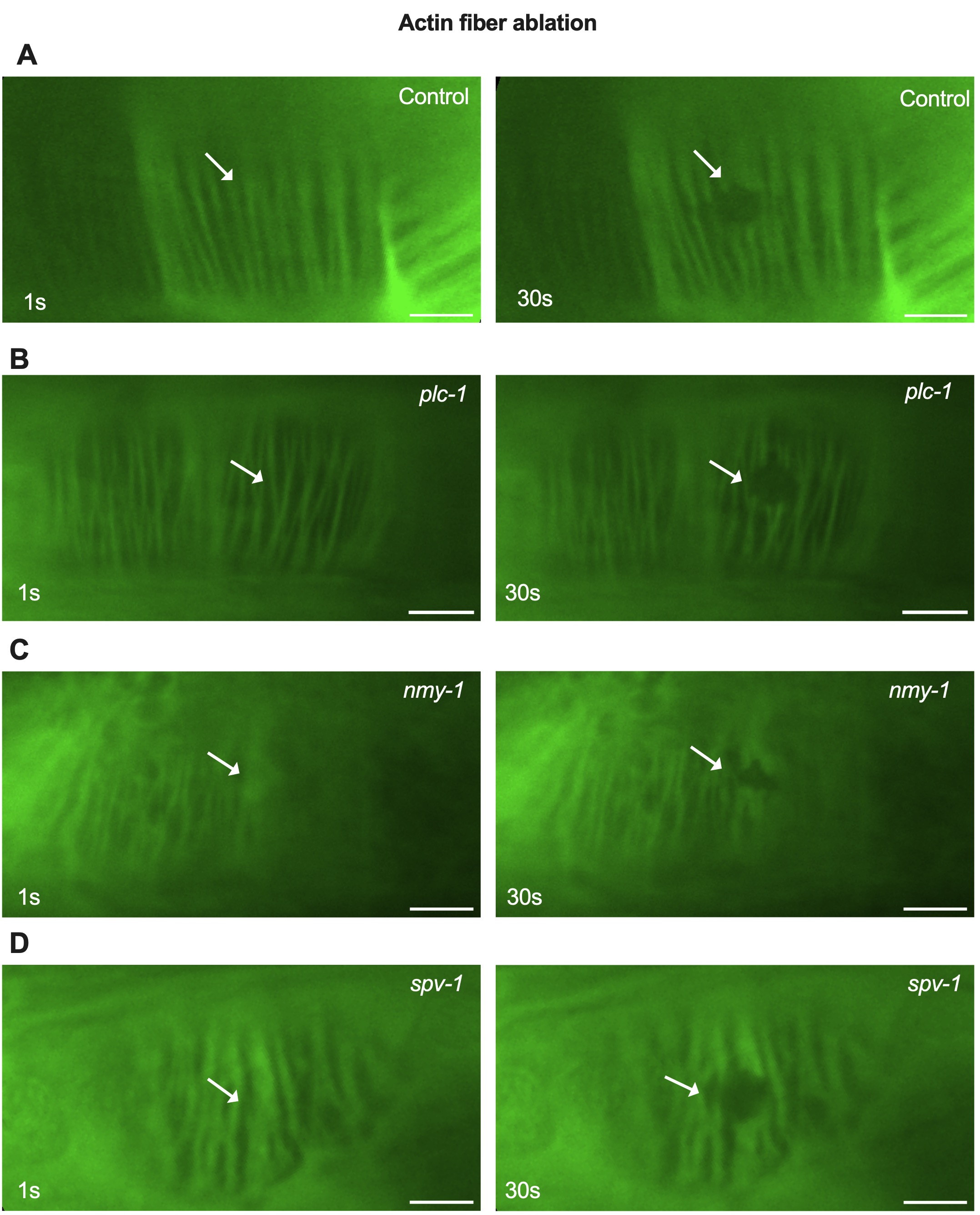

### Supplemental Figure 2

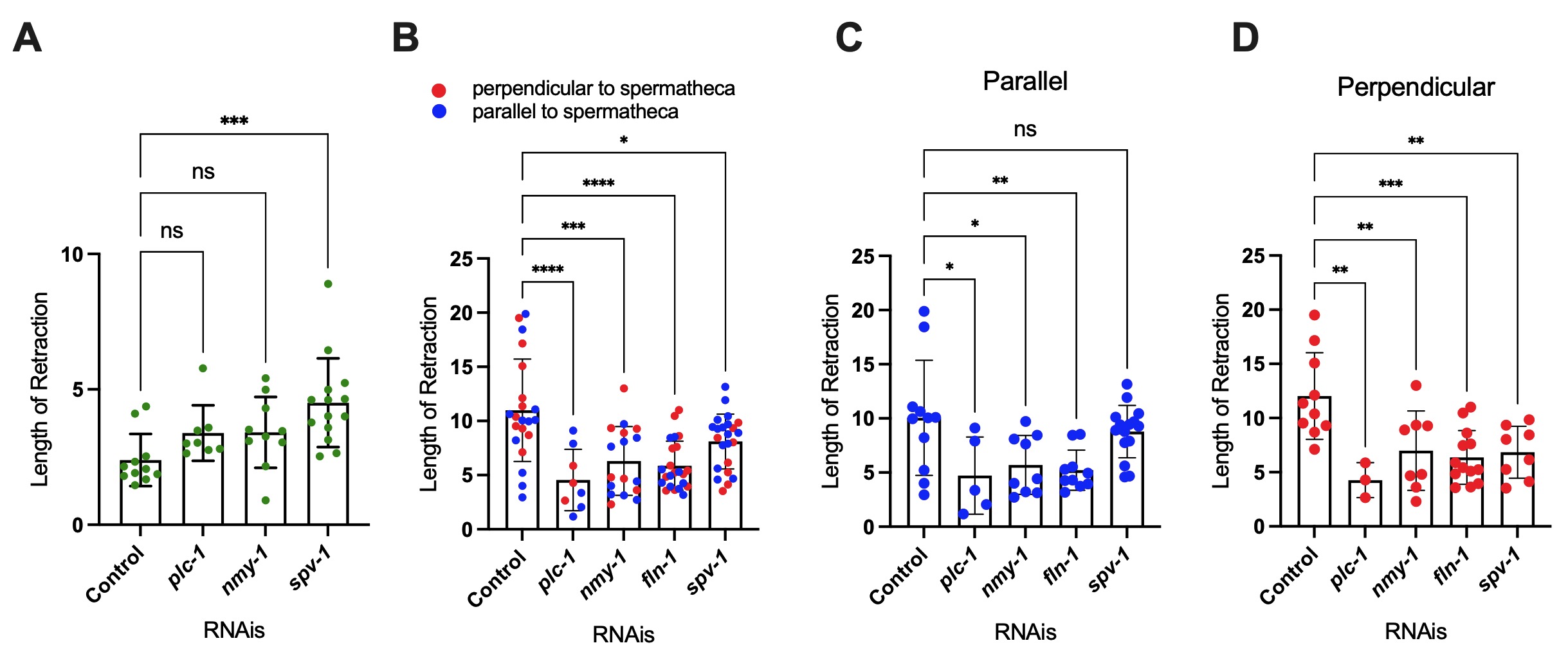

### Supplemental Figure 3

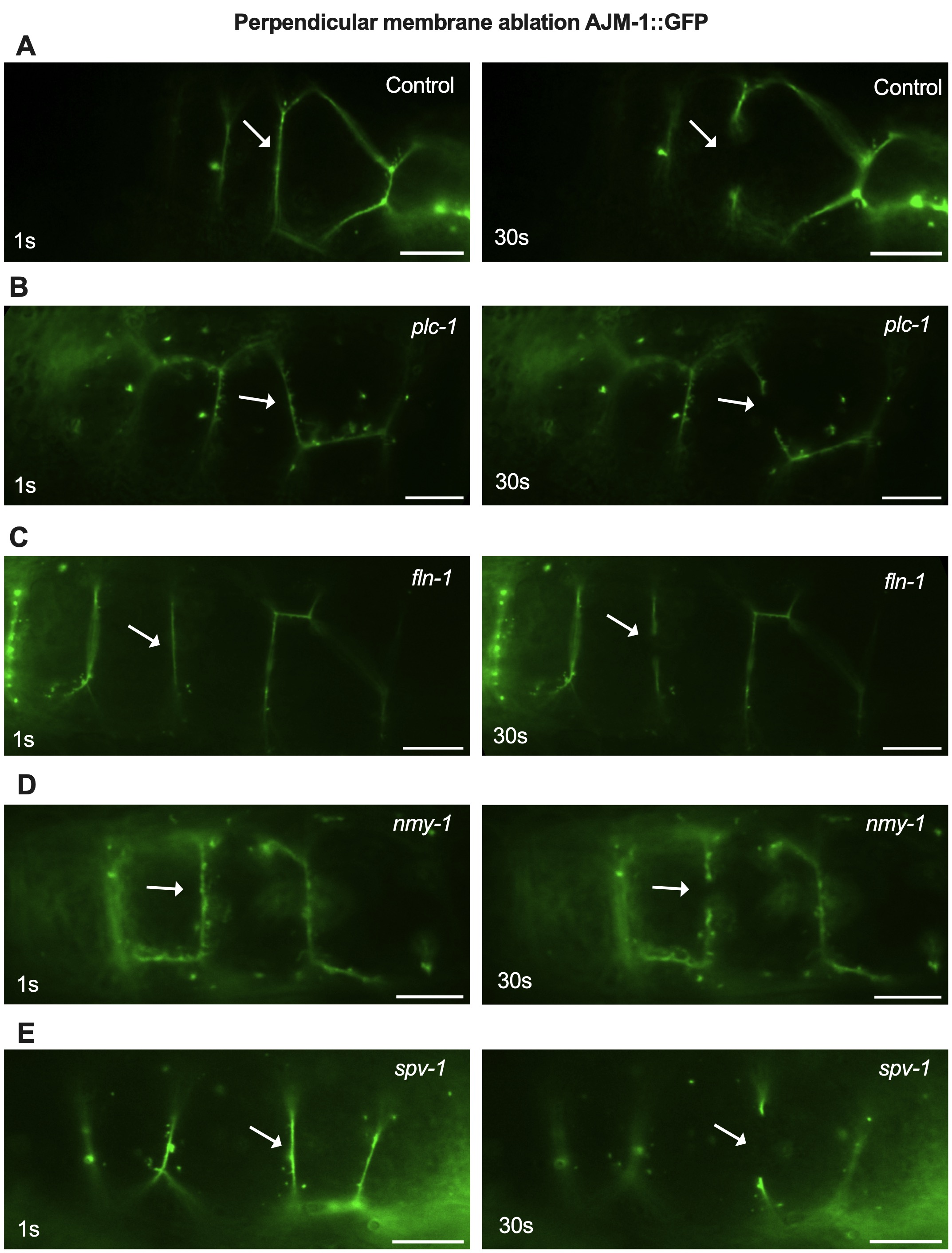

### Supplemental Figure 4

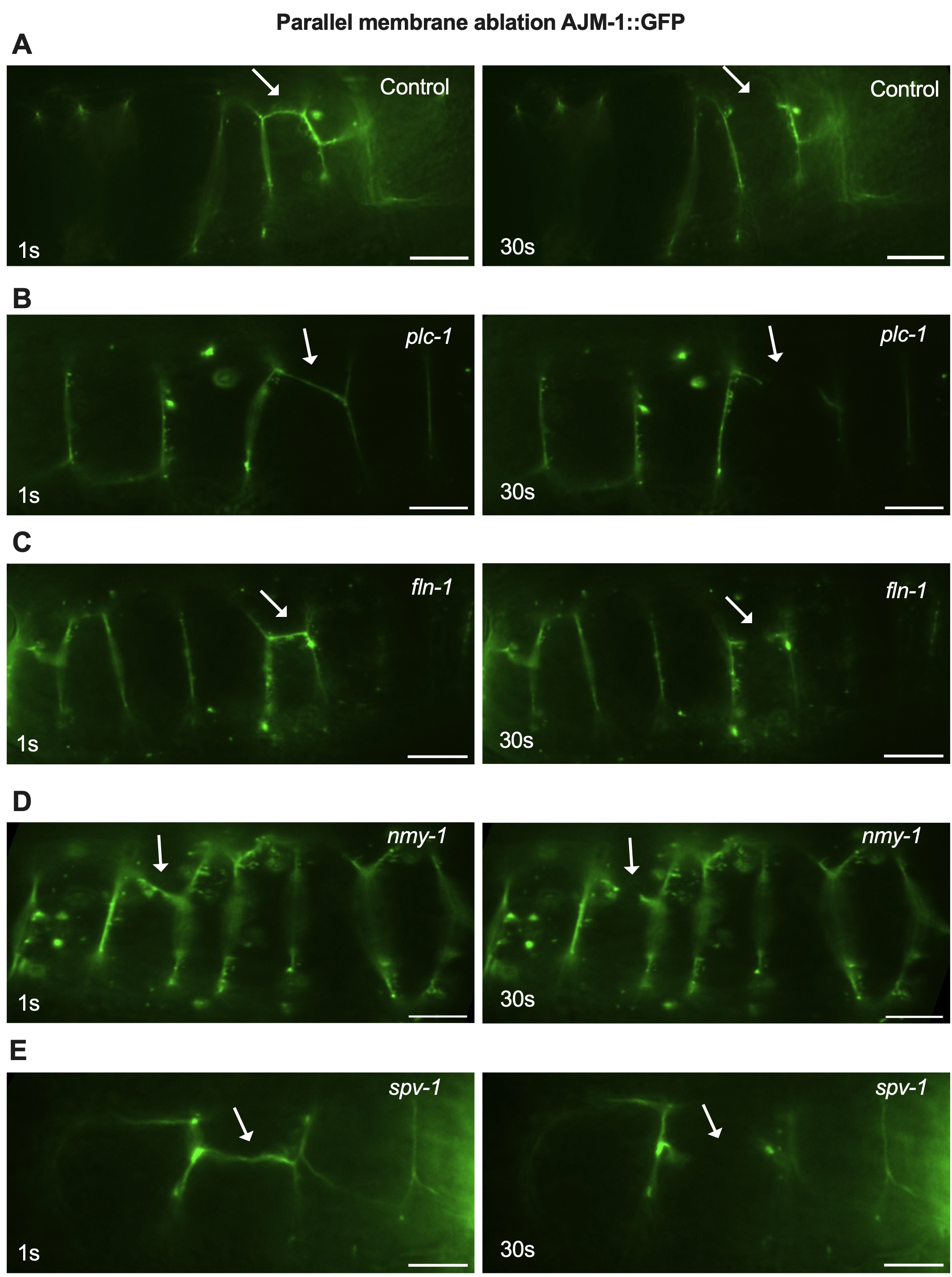
